# CancerDiscover: A configurable pipeline for cancer prediction and biomarker identification using machine learning framework

**DOI:** 10.1101/182998

**Authors:** Akram Mohammed, Greyson Biegert, Jiri Adamec, Tomáš Helikar

## Abstract

**Motivation:** Use of various high-throughput screening techniques has resulted in an abundance of data, whose complete utility is limited by the tools available for processing and analysis. Machine learning holds great potential for deciphering these data in the context of cancer classification and biomarker identification. However, current machine learning tools require manual processing of raw data from various sequencing platforms, which is both tedious and time-consuming. The current classification tools lack flexibility in choosing the best feature selection algorithms from a range of algorithms and most importantly inability to compare various learning algorithms.

**Results:** We developed CancerDiscover, an open-source software pipeline that allows users to efficiently and automatically integrate large high-throughput datasets, preprocess, normalize, and selects best performing features from multiple feature selection algorithms. The pipeline lets users apply various learning algorithms and generates multiple classification models and evaluation reports that distinguish cancer from normal samples, as well as different types and subtypes of cancer.

**Availability and Implementation:** The open source pipeline is freely available for download at https://github.com/HelikarLab/CancerDiscover.

**Supplementary Information:** Please refer to the CancerDiscover README (Supplementary File 1) for detailed instructions on installation and operation of the pipeline. For a list of available feature selection methods, see Supplementary File 2.

## Introduction

Classification of a tissue sample as cancer or normal and among different tissue types facilitates cancer treatment, and machine learning has the potential to improve such classification. High-throughput techniques such as microarrays generate massive amounts of data. One of the challenges of performing cancer classification and biomarker identification tasks using gene expression data from various microarray platforms is to convert the data into machine learning framework readable format. Several machine learning methods and software tools have been developed to study cancer classification (Akay, 2009; Aliferis *et al.*, 2002; Liu *et al.*, 2005; Pirooznia *et al.*, 2008; Mao *et al.*, 2005; Peng, 2006; Duan *et al.*, 2005; Kallio *et al.*, 2011; Kolesnikov *et al.*, 2015; Gao *et al.*, 2013; Zuo *et al.*, 2016). However, different analysis steps have to be performed by various tools, often using different software platforms.

Machine learning tools can be a powerful tool for the analysis of these data. For example, Waikato Environment for Knowledge Analysis (WEKA) (Mark Hall, Eibe Frank, Geoffrey Holmes, Bernhard Pfahringer, Peter Reutemann, 2009; Hall *et al.*, 2009) is a machine learning software environment that serves as a platform for clustering and classification of high-throughput data. However, such platforms require data that have been normalized and otherwise preprocessed to address various technical and statistical challenges such as, expression value differences within the dataset, and differences among microarray plates, for example. Moreover, raw high-throughput data are not organized in such a way that machine learning frameworks can directly process them—normalized data must be formatted into a framework native format (for example, Attribute-Relation File Format ARFF for WEKA). This extensive and manual processing is not only time-consuming but also error-prone, making high-quality, large-scale analyses difficult.

To make cancer classification and cancer biomarker identification more accessible to researchers, we have developed CancerDiscover, a software pipeline, which can, given raw, bulk high-throughput data, normalize the data, generate the ARFF files, and build and evaluate machine learning models for cancer type classification and biomarker identification. Unlike software tools that require manual processing and/or are limited in options such as feature selection and classification algorithms (e.g., GenePattern (Reich *et al.*, 2006), ESVM (Huang and Chang, 2007), Prophet (Medina *et al.*, 2007), Chipster (Kallio *et al.*, 2011)), CancerDiscover is a fully automated pipeline, while providing users with full control over each step. Herein, we describe the software, and demonstrate its utility and flexibility through a case study. We also provide benchmarking statistics for datasets of varying sizes. As a tool for identification of potential biomarkers and drug targets, CancerDiscover is complementary to data repositories and software tools such as Oncomine (Rhodes *et al.*, 2007), INDEED (Zuo *et al.*, 2016) and cBioPortal (Gao *et al.*, 2013) that support advanced data visualization and/or analysis of differential gene expression.

## System and Methods

The presented pipeline consists of existing open source software tools and utilizes publicly available datasets and various performance metrics.

### Data Collection

For the case study, microarray gene expression data was collected from the Broad Institute Cancer Program Legacy Publication Resources database (Cancer Program Legacy Publication Resources) that was first published in Bhattacharjee *et al.* 2001 (Bhattacharjee *et al.*, 2001). The plates used for this article were Human U95A oligonucleotide probe arrays, containing 54,675 probes. GPL96[HG-U133A] Human Genome U133A Array, GPL97 [HG-U133B] Human Genome U133B Array, GPL570[HG-U 133_Plus_2] Human Genome U133 Plus 2.0 Array.

### Affy R package (Normalization, background corrections)

The R module Affy provides an established method for normalization and background correction (Gautier, Cope, Benjamin M Bolstad, *et al.*, 2004). This step is crucial for analyzing large amounts of data which have been compiled from different experimental settings. In this step, individual data files are processed to remove sample bias from the data, which could otherwise introduce a bias in the model. Affy R package provides multiple methods for normalizations and background correction, which can be utilized within CancerDiscover using programmatic flags. For the case study given below, quantile normalization (Bolstad, 2001) and robust multi-chip average (RMA) (Irizarry *et al.*, 2003) were used for normalization and background correction, respectively.

### Machine Learning algorithms and framework

Support Vector Machines (SVMs) and Random Forests were used to construct the models for this study. These machine-learning methods were chosen because of their extensive and successful applications to datasets from genomic and proteomic domains (Statnikov *et al.*, 2008; Mohammed and Guda, 2015). Some of the cancer classification tasks were binary (two classes), and the others were multi-class (more than two classes). Though SVMs are designed for binary classification, they can also be used for multi-class classification by a one-versus-rest approach (Cortes and Vapnik, 1995). The one-versus-rest approach for classification is known to be among the best-performing methods for multi-category classification for microarray gene expression (Statnikov *et al.*, 2005).

Models were also constructed using Random Forests (RF), which can solve multi-category problems natively through direct application. The Random Forests algorithm is well suited to the classification of genomic data because of the following advantages (i) it performs embedded feature selection (ii) it incorporates interactions between predictors: (iii) it allows the algorithm to accurately learn both simple and complex classification functions; (iv) it is applicable to both binary and multi-category classification tasks (Bishop, 2007).

### Performance measure

Accuracy was defined as the overall ability of models to categorize testing sample data correctly. Reported measures included the numbers of *true positives* (TP), *true negatives* (TN), *false positives* (FP), and *false negatives* (FN). A true-positive count is the number of samples in a dataset which were correctly categorized *into* classes. A false-positive count is the number of samples in a dataset which were sorted into the wrong category. A true negative count represents the number of samples which were *not* classified into a class to which they do *not* belong, and false negatives are samples which are *not* classified into the class to which they do *not* belong.

Accuracy, Sensitivity (or Recall), Specificity, and Precision are derived from the measures mentioned above as follows: accuracy is the ratio of correctly predicted samples to the total number of samples. Sensitivity is the proportion of true positives that are predicted as positives. Specificity is the proportion of true negatives which are predicted as negatives, and Precision is the ratio of true positives to the total number of true negatives and true positives. Lastly, F-score is defined as the harmonic mean of Precision and Recall and is calculated by first multiplying precision and recall values, then dividing the resulting value by the total of precision and recall, and finally, multiplying the result by two.

The Accuracy, Sensitivity, Specificity, Precision, and F-Score are given by:

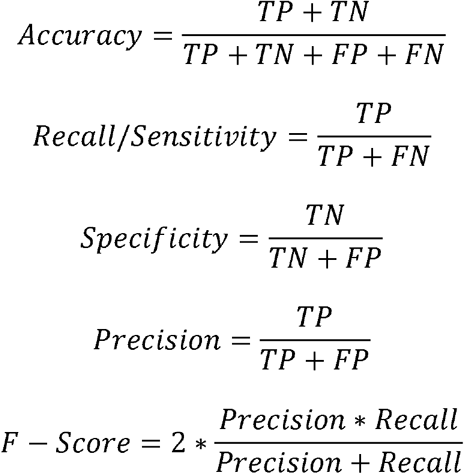

### Model selection and accuracy estimation

For model selection and accuracy estimation, we used 10-fold cross-validation (Mohammed and Guda, 2015; Statnikov *et al.*, 2005). This technique separates data into ten parts and uses nine parts for the model generation while predictions are generated and evaluated by using the one part. This step is subsequently repeated ten times, such that each part (internal test set) is tested against the other nine parts (internal train set). The average performance over the ten accuracies is accepted as an unbiased estimate of the model’s performance.

## Implementation

CancerDiscover is a software pipeline, which takes a raw dataset, normalizes it, generates ARFF files, and builds and accesses machine learning models for cancer type classification and biomarker identification. CancerDiscover consists of eight components: normalization, preliminary feature vector generation, preliminary data partitioning, feature selection, feature vector generation, data partitioning, model training and model testing. These components are organized into three scripts (Normalization, Feature Selection, and Model Building). CancerDiscover utilizes WEKA (Waikato Environment for Knowledge Analysis) (Hall *et al.*, 2009; Mark Hall, Eibe Frank, Geoffrey Holmes, Bernhard Pfahringer, Peter Reutemann, 2009) version 3.8. for its data partitioning, feature selection, and model construction. The pipeline within CancerDiscover is illustrated in Figure 1 and detailed below.

**Fig 1.**
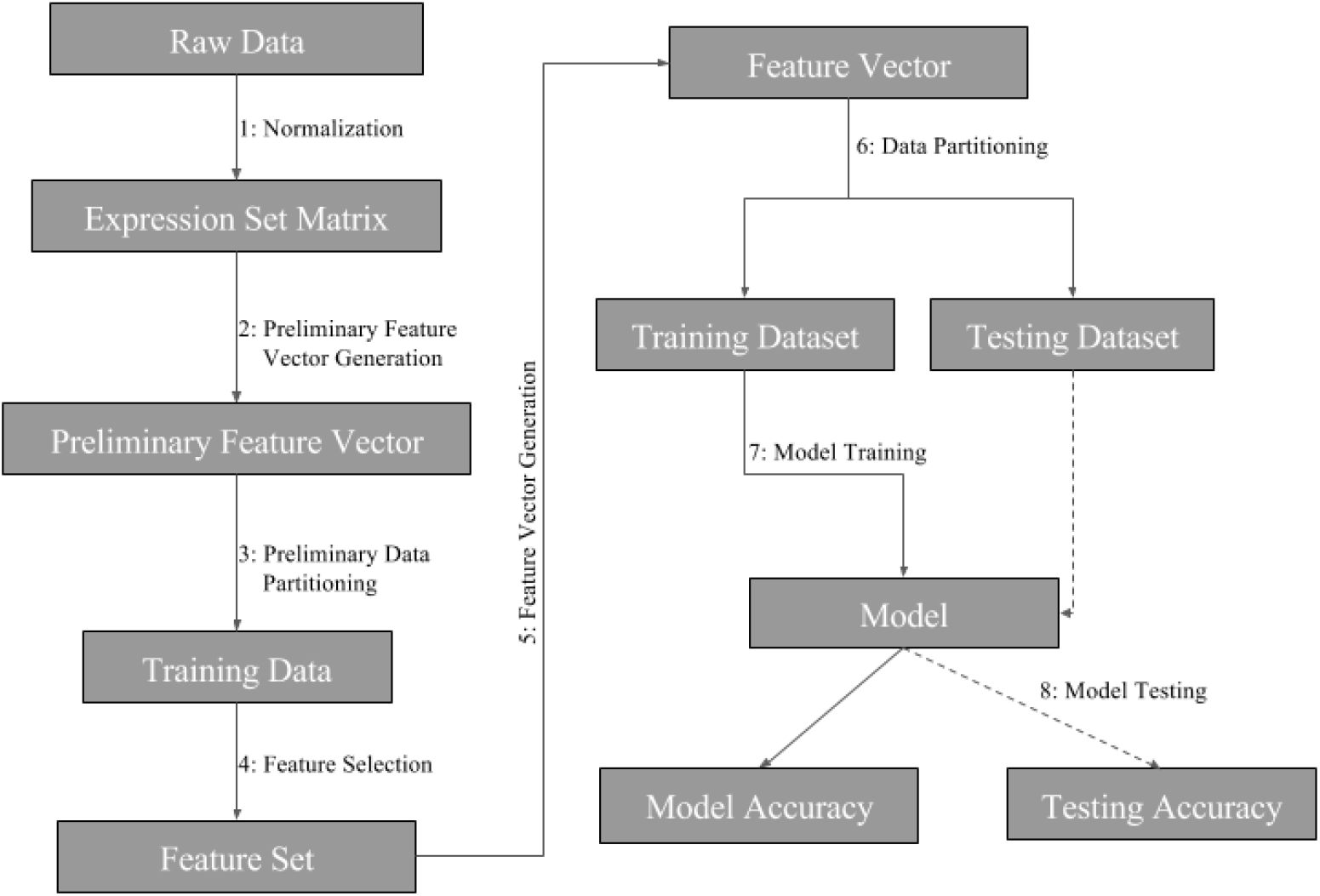
Schematic representation of the CancerDiscover Pipeline: First, raw data are normalized, background correction is performed, and the output is partitioned into training and testing sets. The test set is held in reserve for model testing while the training set undergoes a feature selection method. Feature selection provides a list of ranked attributes that are subsequently used to rebuild the training and testing sets. The training dataset is subsequently used to build machine learning models. Finally, the testing data set is used for model testing.

1. ***Normalization***: Due to the inherent differences among samples obtained from various studies, normalization and background corrections are required to remove or subdue bias in raw data for accurate models. Once raw high-throughput data are obtained, normalization and background corrections are performed using the Affy R module (Gautier, Cope, Benjamin M. Bolstad, *et al.*, 2004) to remove the technical variation from noisy data and background noise from signal intensities and generated the expression set matrix. For the case study, the Quantile Normalization Method (Bolstad and others, 2003) was used to normalize the data, and the background correction was performed using the Robust Multi-Average (RMA) (Irizarry *et al.*, 2003) parameter method by modifying the configuration file.
2. ***Preliminary Feature Vector Generation***: Next, the expression set matrix is used to generate the required ARFF formatted file for WEKA processing. This ARFF file is referred to as the master feature vector.
3. ***Preliminary Data Partitioning (Stratified)***: To maintain an even distribution of sample classes, stratified partitioning has been used by splitting the master feature vector evenly into testing and training sets. These training sets are used to construct the models after feature selection has been performed in the next step. Later, the model’s accuracy will be assessed with the testing set, which had not been exposed to the model, giving an honest assessment of the model. Users of CancerDiscover can specify the size of the data partition of their choice.
4. ***Feature Selection (on training data set only)***: WEKA provides several options for feature selection, and our pipeline allows users to select which algorithms they want to use. Each of these algorithms provides the list of ranked features that distinguish cancer from normal tissue. Once the models with different feature thresholds are built, one can explore the relationship between model accuracy and the number of features considered by the classification algorithms in the evaluation of validation datasets. The feature sets generated are separated into feature thresholds (including the top 1%, 10%, 33%, 66%, 100% of the total number of ranked features as well as the top 25, 50 and 100 ranked features). We arbitrarily chose these thresholds to identify the minimum number of features needed to achieve accurate classification models. For a list of available feature selection methods, see Supplementary File 2.
5. ***Feature Vector Generation***: Since the classification models must be built based only on the ranked features, new feature vectors are generated based on the ranked feature sets discussed in feature selection step.
6. ***Data Partitioning (Stratified)***: Once the new feature vectors (ARFF files) are generated, each feature vectors file will undergo a second data partitioning. This partitioning seed value (or integer that defines the exact sequence of a pseudo-random number) is the same as the one used in the preliminary data partitioning. As such, each new feature vector will be split into the same training and testing sets as in step 3, while the samples used to test the model are avoided for model training. The master training and testing feature vectors and the new training and testing feature vectors differ only in the number of features; the master feature vectors contain all of the features, whereas the newly created feature vectors contain only the features that ranked according to different thresholds.
7. ***Model Training***: WEKA provides various machine learning classification algorithms. Our pipeline is compatible with five diverse classification algorithms and allows the user to build models using only one or all five, as they see fit. The five classification algorithms are Decision Tree (J48) (Iba and Langley, 1992), Naive Bayes (Rish, 2001), IBK (Cover and Hart, 1967), Random Forest (RF) (Breiman, 2001), and Support Vector Machine (SVM) (Cortes and Vapnik, 1995). We used Support Vector Machines (SVM) and Random Forests for the construction of our models in the case study discussed below. These machine learning methods were chosen because of their extensive and successful applications to datasets from both genomic and proteomic domains (Statnikov *et al.*, 2008; Mohammed and Guda, 2015). Each new training dataset informs the machine learning algorithm on model construction using 10-fold crossvalidation. Ten-fold cross-validation separates training data into ten parts and uses nine parts for model generation, while predictions are generated and evaluated using one part. This process is repeatedly performed such that each single tenth of data can be tested against the other nine-tenths of data. After the 10-fold cross-validation, the average performance of all of the folds is used as an unbiased estimate of the performance of model training.
8. ***Model Testing***: Once the models are created, they undergo testing by exposing the model to the companion testing sets that were hidden from the model during model construction. The testing dataset is hidden from the model, such that that the sample classification is based on what the model has learned from the amount of expression in each feature, for every sample, in the training dataset. In the case study below, we illustrate the utility of the software to classify normal vs. cancerous tissues based on gene expression data.

### Installation/operation

All components of the pipeline are organized into three sets of scripts (Normalization, Feature Selection and Model Building and testing), each of which is composed of several scripts (PERL, AWK, SHELL, BASH, R, and SLURM (Simple Linux Utility for Resource Management), installation/operation of the pipeline is described in the Supplementary File 1). SLURM is a computational architecture used to organize user requests into a queue to utilize supercomputer resources. SLURM requires no kernel modifications for its operation and is relatively self-contained. There are two versions of the CancerDiscover pipeline: the beginner version consists of bash scripts that can be run on the local machine, and an advanced version that consists of SLURM scripts that can be run on the supercomputer. Due to the complexity of data manipulation, and/or the sheer size of the high-throughput data, it is recommended to use a supercomputer.

### Case Study

To illustrate possible applications of the pipeline, two kinds of models were generated and tested. The first model was developed to classify tissue samples as either cancerous or normal, according to their gene expression patterns. Sample distributions were as follows: 237 tumor tissue samples and 17 histologically normal tissue samples split evenly into testing and training data sets. Filtered Attribute Evaluator combined with Ranker method was the algorithm selected (using configuration file) to perform feature selection on the training dataset. This algorithm outputs a list of all data features ranked according to their utility in distinguishing the different classes of samples; features ranked at the top of the list are most useful in distinguishing cancer from normal samples. The plates used for this case study contain approximately 10,000 full-length genes corresponding to 54,675 probes (features). Each feature was ranked using the feature selection algorithm, and the top 1%, 10%, 33%, 100% of ranked features as well as additional feature sets containing the top 3, 6, 12, 100, 500, 0.25%, 0.5%, of ranked features were used for generating several models simultaneously. Training and testing accuracies are reported in Figure 2A–D. We selected RF and SVM as the machine learning classification algorithms for this case study.

**Fig 2.**
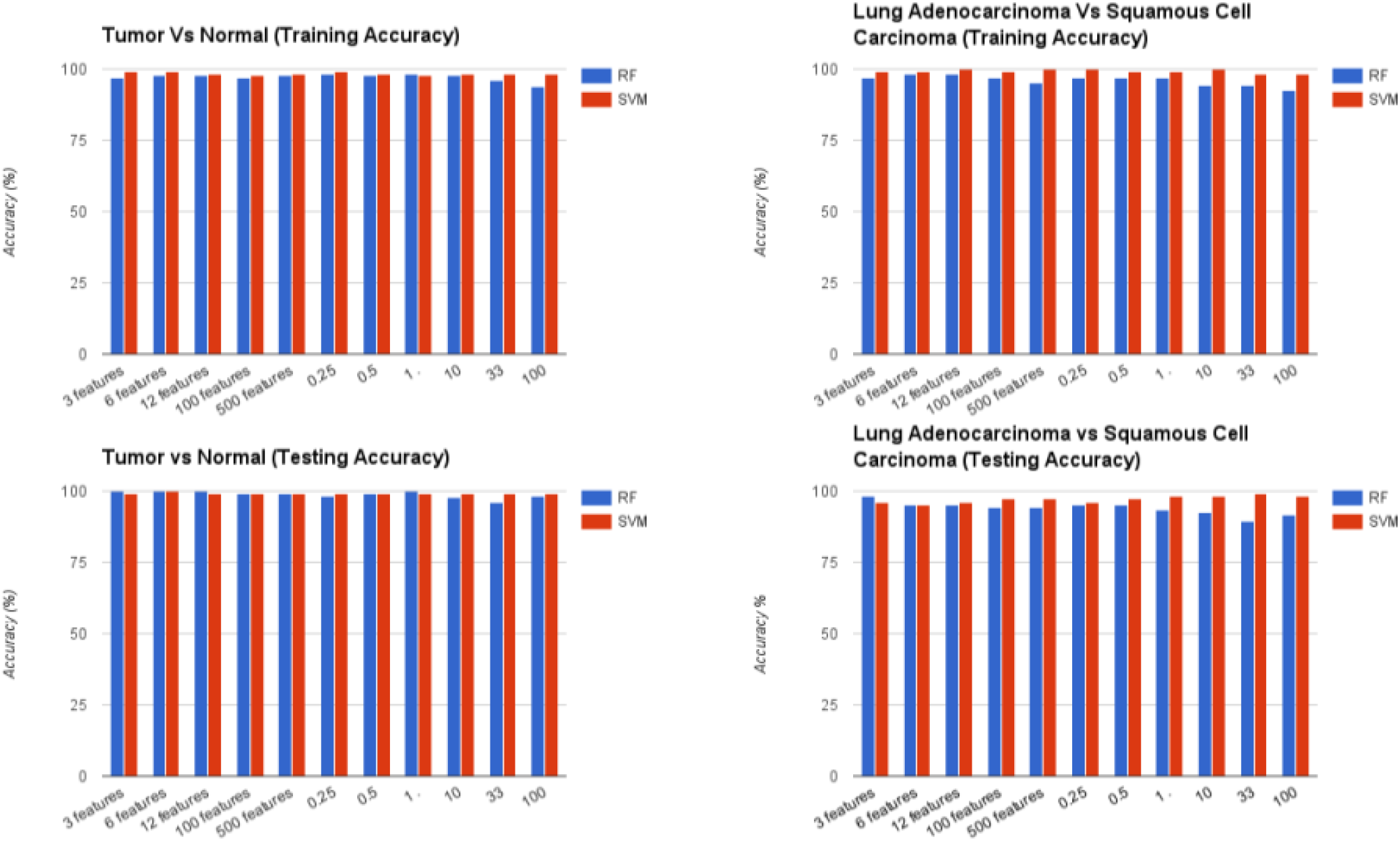
Model Accuracies for the Classification of Tumor vs. Normal and Adenocarcinoma vs. Squamous Cell Carcinoma: RF represents Random Forest classifier and SVM indicates Support Vector Machine classifier. (A, B) Training accuracies for Tumor vs. Normal and Adenocarcinoma vs. Squamous Cell Carcinoma, respectively. (C, D) Testing Accuracies for Tumor vs. Normal and Adenocarcinoma vs. Squamous Cell Carcinoma, respectively.

We achieved a model training accuracy of 98.43% for the RF classifier using the top 0.25% (31 attributes) of features. Models constructed using the top 3% of ranked features reported an accuracy of 96.06%, while using the entire list of features (100%) resulted in the lowest accuracy of 93.70%. Training accuracies for the SVM classifier were 99.21% for the models that used the top 3 features. Accuracy declined with the increasing number of features, with models that used the top 12 ranked features reporting an accuracy of 98.43%. SVM resulted in the lowest (though still relatively high) accuracy of 97.64% using 100 features. This shows that as few as the top 31 features are sufficient to achieve a higher accuracy, using random forest classifiers, whereas top 3 features are sufficient to achieve a higher accuracy using support vector machines.

The second set of models was also bi-class; however, the models were developed to distinguish lung sub-types (adenocarcinoma vs. squamous cell carcinoma), rather than tumor vs. normal tissue. 190 lung adenocarcinoma samples and 21 squamous cell carcinoma samples were evenly split into training and testing datasets. After feature selection, the list of ranked features was used to generate models based on different feature thresholds. Results from testing accuracies can be seen in Figure 2B. With the entire list of ranked features, the RF testing accuracy was 91.51%, increasing in accuracy as the percentage or number of ranked features decreased. The top 1% of ranked features (126 attributes) resulted in model testing accuracy of 93.40% while the top 0.25% (31 attributes) of ranked features resulted in testing accuracies of 95.28%. A similar trend was seen going from the top 500 features to the top 3 features. On average, SVM testing accuracies were more consistent, and higher than those based on RF. The model generated with top 3 features resulted in accuracy of 96.23%, while using the top 6 features resulted in accuracy of 95.28%. Using 100 features resulted in testing accuracy of 97.17%. Using the top 0.25% and 0.5% resulted in accuracies of 96.23% and 97.17%, respectively, while using the top 1% and 10% features resulted in accuracy of 98.11%. Using the top 33% of ranked features resulted in the highest testing accuracy of 99.06%. Confusion matrices for the models generated using the top 3 features are reported in Table 1.

**Table 1:**
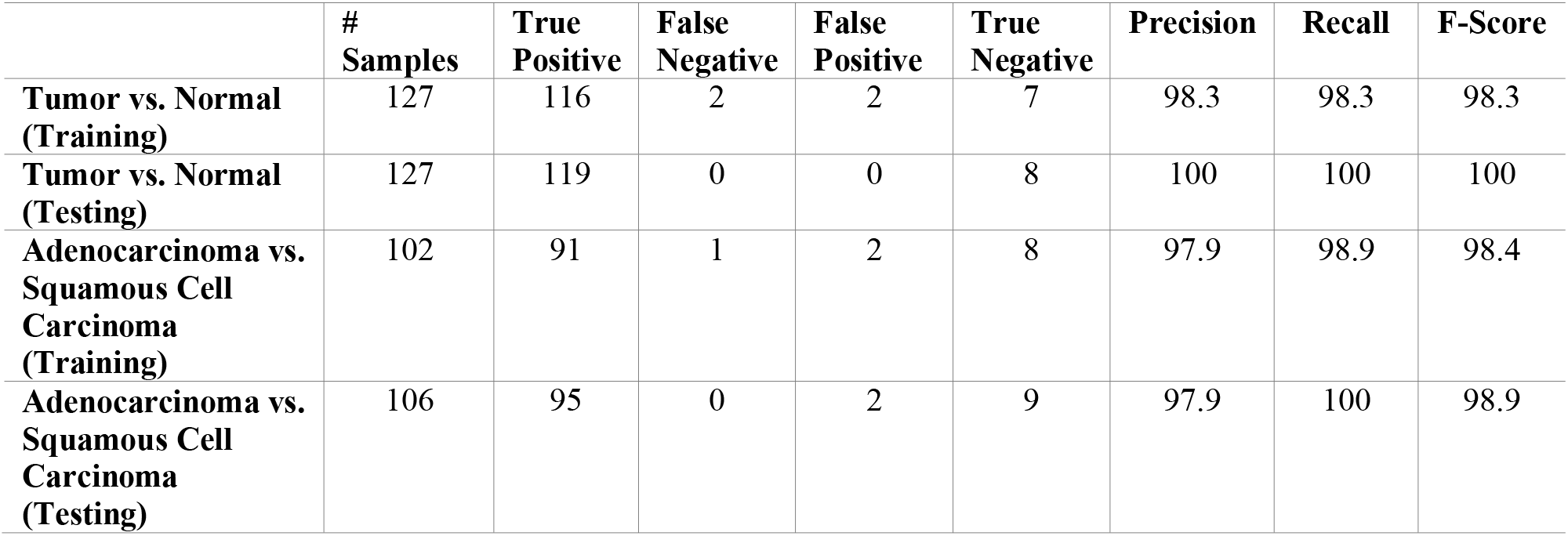
Random Forest Training and Testing Accuracies for Top 3 Ranked FeatureModels.

|  | #<br>Samples | True<br>Positive | False<br>Negative | False<br>Positive | True<br>Negative | Precision | Recall | F-Score |
| --- | --- | --- | --- | --- | --- | --- | --- | --- |
| **Tumor vs. Normal**<br>**(Training)** | 127 | 116 | 2 | 2 | 7 | 98.3 | 98.3 | 98.3 |
| **Tumor vs. Normal**<br>**(Testing)** | 127 | 119 | 0 | 0 | 8 | 100 | 100 | 100 |
| **Adenocarcinoma vs.**<br>**Squamous Cell**<br>**Carcinoma**<br>**(Training)** | 102 | 91 | 1 | 2 | 8 | 97.9 | 98.9 | 98.4 |
| **Adenocarcinoma vs.**<br>**Squamous Cell**<br>**Carcinoma**<br>**(Testing)** | 106 | 95 | 0 | 2 | 9 | 97.9 | 100 | 98.9 |

As shown in Table 1, we were able to achieve a high degree of accuracy using a small fraction of top ranked features (3 features). This case study illustrates the pipeline’s flexibility, utility, and ease-of-use in the generation of several models simultaneously from raw high-throughput data. It also highlights the customization allowed by CancerDiscover on the individual steps of a common high-throughput data analysis pipeline, including the normalization methods, data partitions, feature selection algorithms, classification algorithms, and the threshold or percentage of ranked features for additional analysis.

### CancerDiscover Benchmarking

To assess the performance of the software, benchmarking was performed using acute myeloid leukemia (AML) and normal blood sample expression data downloaded from NCBI (GSE6891, GSE2677, GSE43346, GSE63270) [HG-U133_Plus_2] Human Genome U133 Plus 2.0 Array (Jung *et al.*, 2015; Sato *et al.*, 2013; Schmidt *et al.*, 2006; Verhaak *et al.*, 2009). Samples were collected to perform benchmarking on datasets with specific sample quantities; 500, 200, 100, 50 and 10. Each dataset was run through the pipeline using default settings (perform all possible feature selection and classification algorithms) to determine the required computational resources such as the total amount of elapsed time, the amount of working memory required for each step of the pipeline and the overall total amount of working memory. These factors, mainly depend on the size of the dataset being processed. Benchmarking was performed using computational resources at the Holland Computing Centre of the University of Nebraska which has 106 nodes,4 CPU/ 64 cores and 256GB RAM per nodes. Benchmarking was also performed on a PC (16GB of memory; Intel® Core™ i7-4770 CPU @ 3.40GHz x8) with Ubuntu 64-bit OS. Table 2 below shows the benchmarking results.

**Table 2:**
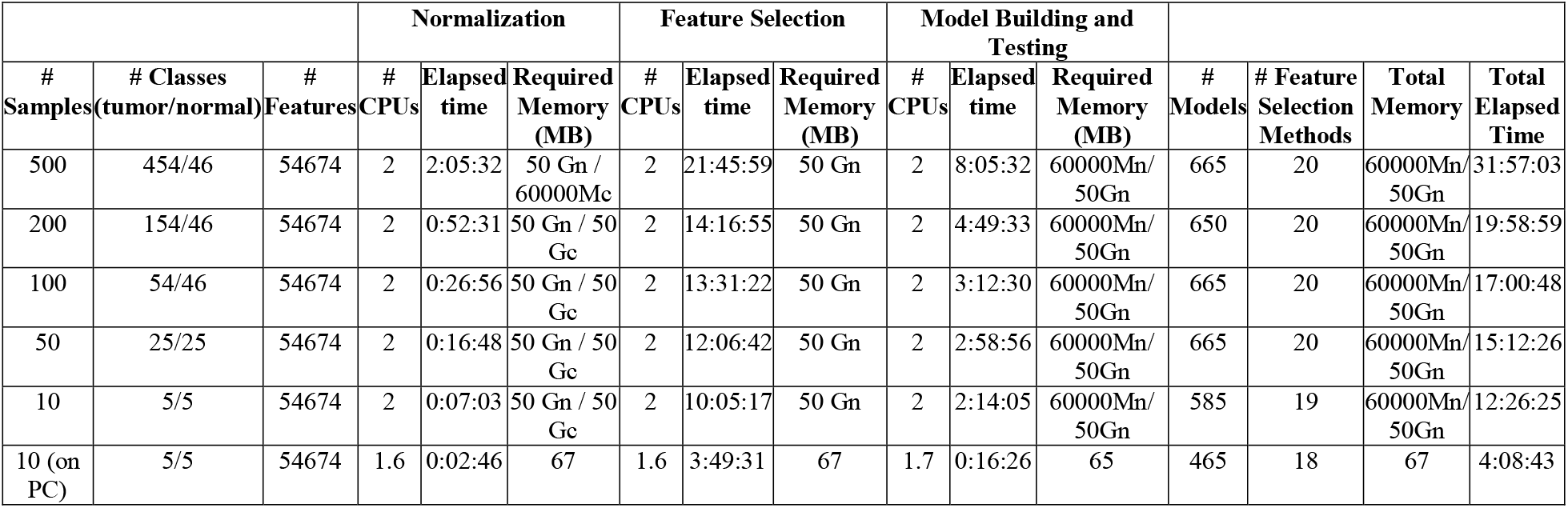
Benchmarking Results. Elapsed time refers to the amount of real-time spent processing that function. CPUs refer to the number of CPUs which were required for that process. CPU Time can then be calculated by multiplying the elapsed time by the number of CPUs.

Starting with the smallest dataset containing only ten samples, of the 23 possible feature selection methods, 19 completed (four feature selection methods could not produce results due to the 10-fold cross-validation). For those 19 feature selection outputs, 585 classification models (some of the ARFF files are empty due to lower threshold selection) were generated. The 50- sample dataset generated 20 out of the 23 possible feature selection results, allowing the next section of the pipeline to generate 665 classification models. When using 100 samples, 20 out of the 23 possible feature sets were produced, and subsequently utilized to generate 665 classification models. The 200-sample dataset provided 20 of the 23 possible feature selection outputs and generated 650 classification models. Lastly, the 500-sample dataset produced 20 out of the possible 23 feature selection outputs, and generated 665 classification models. As the datasets grew, the time required for cancer classification increased linearly (Table 2).

### Comparison of CancerDiscover with other methods and tools

We compared the performance of CancerDiscover with that of three existing methods, GenePattern (Reich *et al.*, 2006) and Chipster (Kallio *et al.*, 2011) and the method described in Aliferis et al. (Aliferis *et al.*, 2003). We used the same train and test datasets to compare the performance CancerDiscover with these methods. Results of this analysis are summarized in Table 3, and discussed in detail below.

**Table 3:** Comparisons of machine learning classification tools and component parts. This table highlights the capabilities of publicly available tools for performing different functions necessary to generate quality models.

| Tool Name | Tumor vs. Normal Accuracy | Adeno vs. Squamous Accuracy | Normalization | background correction | Custom Data Partitioning | Feature Selection |
| --- | --- | --- | --- | --- | --- | --- |
| CancerDiscover | 99.21 | 99.06 | ✓ | ✓ | ✓ | ✓ |
| GenePattern | 98.43 | 99.06 | ✓ | - | - | - |
| Chipster | 97.63 | 98.82 | ✓ | - | - | - |
| Aliferis et al. | 94.97 | 96.83 | ✓ | ✓ | ✓ | ✓ |

GenePattern (Reich *et al.*, 2006) is a web-platform that allows users to upload data in various forms and performs statistical analysis, class prediction, and classification. Due to the nature of the data used in this study, only SVM classification suite was used to draw comparisons between CancerDiscover and GenePattern. Because GenePattern could not perform normalization and background correction for the given datasets, we used the data normalized by the CancerDiscover pipeline (using RMA method) and provided the normalized data to the SVM classification module of GenePattern. The input data contained all probes (as GenePattern does not provide feature selection options). Machine learning classification models were generated using the training data with accuracies of 98.43% for the Tumor vs. Normal model, and 99.06% for the Adenocarcinoma vs. Squamous Cell Carcinoma model. These higher accuracies could also be due to the normalization and background correction performed by the CancerDiscover.

Of all the three compared software tools, GenePattern’s accuracies are most similar to the ones produced by CancerDiscover – 99.21% and 99.06%, respectively. All probes were utilized in the model building since feature selection could not be performed using GenePattern. On the other hand, CancerDiscover was able to make similarly accurate predictions using as few as three probes (See Table 3). Finally, CancerDiscover differs from the proprietary GenePattern by the fact that CancerDiscover is open-source; as such, its methodologies are transparent and reproducible, and the community can further expand the software.

Chipster is developed based on a client-server architecture. Data is imported at the client side, while all processing is performed on the server side using R. It requires that all data needs to be transferred between client and server for each analysis step which can be very time-consuming if the datasets are enormous (Koschmieder *et al.*, 2012). Chipster was not able to successfully perform a classification when we provided the dataset containing all probes. As a result, feature selection was performed artificially; that is, the datasets provided to Chipster contained only those probes selected by our CancerDiscover feature selection method; thus datasets provided contained the top 3, 6, 12, 100, or 500 probes. Raw data in the form of CEL files were normalized (RMA normalization) by Chipster. The accuracy using top 3 probes for the Tumor vs. Normal model was 97.63%, whereas, for the Adenocarcinoma vs. Squamous Cell Carcinoma model was 98.82%, ranking 3^rd^ for the accuracy assessment (Table 3). These accuracy assessments for the CancerDiscover are better than the results provided by Chipster.

Data used in this paper were also analyzed independently in Aliferis et al. (Aliferis *et al.*, 2003), using two feature selection algorithms: Recursive Feature Elimination and Univariate Association Filtering. These algorithms identified 6 and 100 features, respectively, as significant for cancer vs. normal classification, and 12 and 500 features, respectively, for adenocarcinoma vs. squamous cell carcinoma classification. Aliferis et al. reported average accuracies across classification algorithms: 94.97% for cancer vs. normal model, and 96.83% for the squamous carcinoma vs. adenocarcinoma model. In comparison, CancerDiscover resulted in 99.21% accuracy for cancer vs. normal model, and 99.06% for the adenocarcinoma vs. squamous cell carcinoma model, while using only three features. In the context of these data, CancerDiscover was more accurate, while using less information than that of Aliferis et al.

These results demonstrate that the CancerDiscover method is complementary to some of the existing methods, such as GenePattern, Chipster, and Aliferis et al. methods, and that it is also suitable for accurate classification of other types of cancer types and subtypes. Although the classification accuracy of CancerDiscover was higher than that of the compared methods, the strengths of CancerDiscover lie in its streamlined nature that enables users to begin with raw data and proceed to assessable machine learning models within a complete pipeline. Another strength of CancerDiscover is that it is flexible, allowing users to utilize various methodologies within the platform, and further extend the software as a whole due to its open-source nature.

## Conclusion

We have developed a comprehensive pipeline, CancerDiscover, which enables researchers to automate the processing and analysis of high-throughput data with the objective of classifying cancer and normal tissue samples (including cancer sub-types). Herein, we showcased the pipeline’s flexibility, utility, and ease-of-use in generating several models simultaneously from raw data. CancerDiscover allows users to customize each step of the pipeline, selecting individual normalization methods, data partitions, feature selection algorithms, and classification algorithms for additional analysis. The CancerDiscover pipeline was able to accurately classify the sample data using an optimal number of top ranked features. Although we only discussed binary models here, multi-class models can also be generated and validated. Benchmarking demonstrated the high performance of the pipeline across datasets of varying sizes. Although we used a dataset as small as ten samples, larger datasets provide more information for machine learning algorithms to accurately classify samples. Researchers who might have hundreds of samples can now utilize machine learning tools for diverse projects, including biomarker identification, drug response, and tissue classification without extensive technical knowledge, while retaining significant flexibility.

## Author Contributions

TH and JA conceptualized the project. AM and GB wrote the software and developed the case study, under the guidance of AM. All authors wrote, reviewed, and revised the manuscript.

## Funding Information

Supported by National Institutes of Health grant # [1R35GM119770- 01 to TH].

## Competing interests

The authors have no competing interests.

## Acknowledgments

We would like to thank Lara Appleby for comments on previous versions of this manuscript.

